# SAP domain facilitates efficient loading of Ku onto DNA ends

**DOI:** 10.1101/2023.06.26.546499

**Authors:** Jaroslav Fulneček, Eva Klimentová, Albert Cairo, Sona Valuchova Bukovcakova, Panagiotis Alexiou, Zbynek Prokop, Karel Riha

## Abstract

Recognition and processing of DNA ends play a central role in maintaining genome integrity. The evolutionarily conserved DNA repair complex Ku serves as the primary sensor of free DNA ends in eukaryotic cells. Its rapid association with DNA ends is crucial for several cellular processes, including non-homologous end joining (NHEJ) DNA repair and telomere protection. In this study, we conducted a transient kinetic analysis to investigate the impact of the SAP domain on individual phases of the Ku-DNA interaction. Specifically, we examined the initial binding, the subsequent docking of Ku onto DNA, and the sliding of Ku along DNA. Our findings revealed that the C-terminal domain of Ku70, known as SAP ((SAF-A/B, Acinus and PIAS), facilitates the initial phases of Ku-DNA interaction, but does not affect the sliding process. This suggests that SAP may either establish the first interactions with DNA, or stabilize these initial interactions during loading. To assess the biological role of SAP, we generated Arabidopsis plants expressing Ku lacking the SAP domain (ΔSAP). Intriguingly, despite the decreased efficiency of the ΔSAP Ku complex in loading onto DNA, the mutant plants exhibited full proficiency in classical NHEJ and telomere maintenance. This indicates that the speed of Ku loading onto telomeres or DNA double-strand breaks (DSBs) is not the decisive factor in stabilizing these DNA structures.

## Introduction

The DNA repair complex known as Ku plays a central role in maintaining genome integrity. Ku is composed of two subunits, Ku70 and Ku80, forming a heterodimer that binds to DNA ends. It acts as an initial responder to DNA damage, rapidly recognizing and binding to DNA double-strand breaks (DSBs) (1,2). Once bound, Ku stabilizes DSBs by preventing their degradation, and recruits and activates other DNA repair proteins, initiating a cascade of DNA damage signaling and processing events that culminate in the rejoining of broken DNA ends. This DNA repair mechanism is known as non-homologous end joining (NHEJ) and represents the major DSB repair pathway in the G1/G0 phase of the cell cycle. Ku is evolutionarily conserved across all eukaryotes and it is the defining component of the canonical c-NHEJ. When c-NHEJ is impaired, DSBs can be repaired by alternative end-joining pathways (alt-EJ) that may result in more extensive resection of DNA ends, and their lower accuracy increases the risk of chromosomal rearrangements (3).

In addition to its role in c-NHEJ, Ku has also been implicated in other aspects of DNA end metabolism including detection of DNA viruses (4), V(D)J recombination, and stabilization and restart of arrested replication forks (5,6). Furthermore, Ku is important for maintenance and protection of telomeres, specialized chromatin structures that shield natural ends of linear eukaryotic chromosomes from being recognized as DSBs (7,8). Ku-deficient organisms often exhibit chromosome end deprotection characterized by nucleolytic resection of telomeres, increased recombination and chromosome fusions. The involvement of Ku at telomeres is seemingly contra-intuitive, because one of the key hallmarks of telomere dysfunction, chromosome end-to-end fusions, are mediated by NHEJ. Thus, it is assumed that Ku activity is modulated in the context of telomeres to assure end protection while preventing downstream steps of NHEJ.

These functions of Ku are implemented through Ku binding to free DNA ends. Ku exhibits a strong sequence-independent affinity to DNA ends and is assumed to be the major DNA-end binding factor in eukaryotic cells (8). The affinity to DNA ends stems from the structural properties of the Ku heterodimer. The Ku70 and Ku80 proteins share a similar topology, consisting of an N-terminal α/β domain, a central antiparallel β-barrel, and C-termini composed of subunit-specific globular domain connected with the protein core by a flexible linker. The Ku heterodimer forms a ring-like structure with a positively charged channel through which DNA is threaded, resembling a bolt and nut mechanism. (9). Ku is loaded onto DNA directionally with the Ku80 side of the heterodimer oriented towards the DNA end (10). Mutations altering the electrostatic charge in the leading part of the channel abolish Ku-DNA interaction (11,12), validating the essential role of the channel in DNA binding. Once loaded onto DNA, Ku can freely slide along duplex DNA using an energy-free mechanism that is not fully understood.

Another part of Ku implicated in DNA interaction is the C-terminal region of Ku70, which includes a flexible loop and the SAP (SAF-A/B, Acinus and PIAS) domain. The SAP domain was identified through a bioinformatic search as a putative DNA binding motif that is enriched in proteins involved in chromosomal organization, DNA repair, and transcription (13). The Ku70 SAP domain adopts a unique helix-extended loop-helix structure with patches of positively charged residues on its surface, suggesting potential interaction with DNA (9,14). Indeed, the SAP domain has been reported to interact with DNA, although at a much lower affinity than the Ku70/80 central channel, and deletions spanning the Ku70 C-terminus have been shown to impair Ku-DNA binding in vitro (14–16). Cryo-EM studies indicate that the SAP domain does not assume a fixed position in the complex but undergoes a change in its location upon DNA binding (17,18). Interestingly, the SAP domain in the DNA-bound Ku complex is positioned distally from the central channel and does not directly contact DNA, raising a question of how SAP contributes to DNA binding. Furthermore, the contribution of the SAP domain to DNA repair and genome stability has not yet been assessed. Thus, its precise role of SAP in Ku-DNA interaction and its biological functions still remain to be resolved.

Plants lack DNA-PKcs, but they possess the core NHEJ proteins including the Ku complex suggesting that mechanism of plant NHEJ is analogous to yeast ad mammals (19). Ku is dispensable in Arabidopsis, and ku mutants, except for increased sensitivity to genotoxic stress, do not exhibit any discernable growth or developmental defects (20–22). Inactivation of Ku also leads to partial telomere deprotection characterized by telomerase-mediated telomere extension, exonucleolytic resection of chromosome ends, and increased telomeric recombination (23–25). Protection of Arabidopsis telomeres is mediated by physical association of Ku with chromosome termini through the central DNA binding channel, which parallels its association in the context of DSBs (12). However, requirements for Ku-DNA binding differ at telomeres and in NHEJ. Mutations in the central channel of the Ku complex, impairing its translocation along DNA, preserve telomere protection but result in deficient DNA repair (12). These observations suggest that in the context of a DSB, Ku translocation is required to free the end for downstream processing and ligation, while entrapment of Ku at chromosome termini is sufficient for telomere protection.

The ability to functionally distinguish the Ku-DNA binding requirements in two biological processes prompted us to examine the role of the Ku70 SAP domain in Arabidopsis. In this study, we analyzed effect of the SAP domain on kinetics of Ku-DNA interaction in vitro, and examined contribution SAP to c-NHEJ and telomere maintenance in vivo.

## Material and Methods

### Plant material

The Arabidopsis thaliana ecotype Col-0 ku70-2 line (SALK_123114) was obtained from the Nottingham Arabidopsis Stock Centre (NASC). Complementation vectors and transgenic plants At18 and At25 were previously described (12). Plants were grown in phytotrons under LED illumination (white 77% /red 20% /infrared 3%; 150 μmol m-2 s -1) with 16/8 h light/dark regime. Genotyping was carried out by PCR with primers indicated in Table S1.

### Plant transformation

Plant binary vector for production of ΔSAP plants was derived from pCBSV70wt that encoded the full-length Ku70 (12) by deleting nucleotides coding for L593-K621 using in vitro mutagenesis based method (26) with primers listed in Table S1. Binary vectors were electroporated into Agrobacterium tumefaciens GV3101 and Arabidopsis plants heterozygous for ku70-2 were transformed by the floral dip method. Transformed plants were selected on soil sprayed with 40 μg/ml BASTA. ku70 mutants homozygous for the transgene were used for analysis. Protein levels in transgenic plants were examined by immunoblotting using rabbit anti AtKu70 (12) and detected on LiCor Odyssey using goat IRDye 800 CW α- rabbit (LI-COR Biosciences).

### Telomere analyses

Terminal restriction fragment analysis and t-circle amplification were performed as previously described (23,25).

### Genotoxicity assays

Arabidopsis seeds were surface sterilized by Cl_2_ evaporation, and approximately 50 seeds were put on 0.5x Murashige and Skoog (MS) agar plates supplemented with the concentration range of bleomycin (Calbiochem). Seeds were stratified at +4°C for two days and then incubated at 21°C in cultivation chambers (16/8 hr light/dark cycle, light 60 μmol m-2 s -1). Seedlings were scored 11 days after germination. Root growth assay was performed in a similar way in square plates positioned vertically in holders at angle of 80° degrees.

### Protein expression and purification

The expression vector pFastBac Dual (Invitrogen) was used to co-express Arabidopsis Ku70 and His-tagged Ku80 subunits in insect cells using a baculovirus expression system at Vienna BioCenter Core Facility as described previously (12). Vector for production of ΔSAP Ku70 was generated by deleting nucleotides coding for L593-K621 from pFastBac Dual Atwt plasmid (12) using in vitro mutagenesis-based method. Briefly, the pFastBac Dual-based plasmids were transposed to EMBacY bacmid in E. coli and subsequently transfected to Sf9 cells for baculovirus generation. The cells were infected with virus and harvested on DPA3. Cells were resuspended in lysis buffer (50 mM Tris-Cl, 250 mM KCl, 10% v/v glycerol, 1 mM DTT, pH 8.0) supplemented with protease inhibitors (Roche) and frozen in liquid nitrogen. Thawed cells were spun and Ku was bound to His Mag Sepharose Ni (GE Healthcare). Beads were washed in Lysis buffer containing 50 mM imidazole and Ku was eluted in Lysis buffer with 250 mM imidazole. Proteins were filtered using Nanosep centrifugal columns (Pall) and stored on ice.

### DNA binding assays

EMSA was performed as previously described (12). mwPIFE was measured in Pierce NeutrAvidin coated black 96well plates (#15117, Thermo Scientific) in duplicate according published protocol (27). PIFE was measured in 50 µl of total volume with one pmol of immobilized Cy-3 labeled dsDNA probes and 6 pmol of Ku complex. Cy3-labelled DNA probes were prepared by annealing synthetized DNA oligonucleotides in annealing buffer (10mM TRIS-Cl pH=7.5, 50mM NaCl, 1mM EDTA) by heating to 95°C for 1 min in ThermoMixer C (Eppendorf) and cooling down to room temperature. Oligonucleotides used for generating the probes are listed in Table S1. Stop-flow PIFE was measured using a three-syringe SFM-300 device equipped with a microcuvette µFC 08 cell (8 µl) and combined with a manual monochromator spectrometer MOS-200 equipped with a Xe arc 19 lamp (BioLogic, France). The instrument was operated by a BioKine 32 v 4.63. We applied excitation light at 547 nm and detected emission using a 585/65 ET Bandpass filter (AHF Analysentechnik) in 1 ms intervals for 1 s and 10 ms intervals for 20 s. The measurements were performed in the following buffer (150 nM KCl, 35 mM TRIS-Cl pH 8.15, 1 mM EDTA, 0.1 mM DTT, 3.893 mM imidazole and 5.156% glycerol). 250 nM protein solution (the first syringe) was premixed with buffer (the second syringe) in a ratio from 0:10 to 10:0 and the premix was then mixed with 25 nM DNA oligonucleotide and, in the case of 75 bp probe, also with 250 nM NeutrAvidin (Thermo Scientific, the third syringe) in a ratio 1:1. An average trace was composed of at least five traces and was used for analysis. The fluorescence traces were fitted to single or double exponentials using KinTek Global Kinetic Explorer version 5.2 (KinTek, USA) providing the observed rate constants and amplitudes of individual kinetic phases (28).

### DNA end joining assay

We used a 3400 bp DNA fragment (Figure S4) containing 35S-EYFP gene derived from pGWB442 as a substrate for the end joining assay. The fragment was cleaved out from pUC19 by SmaI, purified through agarose gel electrophoresis using NucleoSpin Gel and PCR Cleanup (Macherey-Nagel), eluted in water and concentrated SpeedVac Concentrator SAVANT SPD 121P (Thermo Fisher Scientific). The fragment was transfected in Arabidopsis mesophyll protoplasts according to (29) as follows: protoplasts were isolated from leaves of 4-week-old Arabidopsis grown on soil in a growth chamber at 22°C under 12 /12 h light/dark cycles. 20-30 leaves were cut using a razor blade and digested in 15 ml of digestion solution (1% cellulase Onozuka R10 [Duchefa], macerozyme R10 [Duchefa], 0,4 M mannitol, 20 mM KCl, 20 mM MES pH=5.7), first for 20 min in vacuum followed by 3 h in dark at room temperature. The released protoplasts were filtered with a 70 µm mesh, washed twice with W5 medium (154 mM NaCl, 125 mM CaCl2, 5 mM KCl, 2 mM MES pH=5.7) and stored on ice. For transfection, the protoplasts were resuspended in MMg solution (0,4 mM mannitol, 15 mM MgCL2, 4 mM MES pH=5.7) at 3x10^5^ cells/ml. 100 µl of protoplasts were mixed with 10 µg of linearized DNA in maximally 10 µl volume and 110 µl of PEG solution (4 g PEG 4000, 2.5 ml mannitol 0.8M, 1 ml CaCl2 1M, 3 ml H20) and incubated 10 min in the dark at RT. 440 µl of W5 medium was added to the mixture and, after centrifugation at 700 g for 3 min, the protoplasts were resuspended and incubated in 1 ml of fresh W5 medium for 16 h in the dark at room temperature. Transfected protoplasts were imaged for GFP fluorescence using a Zeiss LSM780 confocal microscope (objective LCI Plan-Neofluar 63x/1.3 1mm Korr DIC M27), to assess transfection efficiency. Next, the protoplasts were collected for DNA extraction. After centrifugation at 700 g for 3 min, the supernatant was discarded and the pellet was frozen at -20 °C for further extraction with the DNeasy® Plant Mini Kit (Qiagen). Total DNA was eluted in 10 µl of water and concentration was measured using NanoDrop. Re-joined ends were amplified by 35 cycles of PCR from 0.8 µl template using Q5 High-Fidelity DNA Polymerase (New England Biolabs) and primers NOSprom-nested-F2 and TagRFP-nested-R2 (Table S1) in 20 µl reaction volume. Five tubes of each amplicon were pooled and DNA purified using NucleoSpin Gel and PCR Clean-Up (Macherey-Nagel). DNA was eluted in 20 µl of Elution Buffer and concentration was measured using Qubit 3.0 fluorometer (Life Technologies).

Sequencing libraries were prepared using KAPA Hyper Prep kit (Roche), using 1000 ng of amplicon DNA according to the manufacturer’s protocol. We omitted any fragmentation steps and size-selected final libraries with AmpureXP magnetic beads (Beckman Coulter). The library was not amplified in the final step of preparation. Libraries were sequenced using the Illumina NextSeq instrument in mid output 2x150 cycles paired-end mode.

Raw FASTQ files from sequencing were quality checked, adapters and low-quality reads were trimmed using Trim Galore (v0.6.7) (https://github.com/FelixKrueger/TrimGalore) and reads shorter than 20 nt were discarded. The paired-end reads were merged with PEAR (v0.9.6) (30). The pre- processed reads were subsequently mapped to the custom reference sequence with STAR (v2.7.10a) (31). Mapped reads were additionally post-processed to be able to count and visualize the different insertion and deletion types around the breakpoint. The reads were filtered to gain only reads containing insertion or deletion and clustered based on their mapping profile in the reference. The final file with clustered reads was visualized in IGV (32). The pipeline used for the processing from sequenced reads to the generation of visualizations is available at https://github.com/evaklimentova/DSB_pipeline. At least 50% of individual clusters per sample were manually evaluated for position and the length of deletion and quantified.

## Results

The Ku70 SAP domain is evolutionarily highly conserved. It is present in plants, animals and fungi (Figure S1), suggesting its origin at the root of eucaryotic life. Arabidopsis SAP domain shares 38% similarity with human SAP and contains nuclear localization signal in its N-proximal part (Figure 1A). The Ku structure has so far been experimentally resolved only in humans. Nevertheless, AlphaFold prediction indicates that Arabidopsis Ku heterodimer forms an asymmetric ring-like structure highly similar to the crystal structure of human Ku (Figure 1B) (9). The SAP domain of Arabidopsis Ku70 is linked to the core of the complex with a long flexible linker and the AphaFold predicted helix- extended loop-helix structure perfectly overlays with the NMR-solved structure of human SAP (Figure 1C) (14). Thus, Arabidopsis Ku appears to have very similar topology and structure like human Ku.

**Figure 1.**
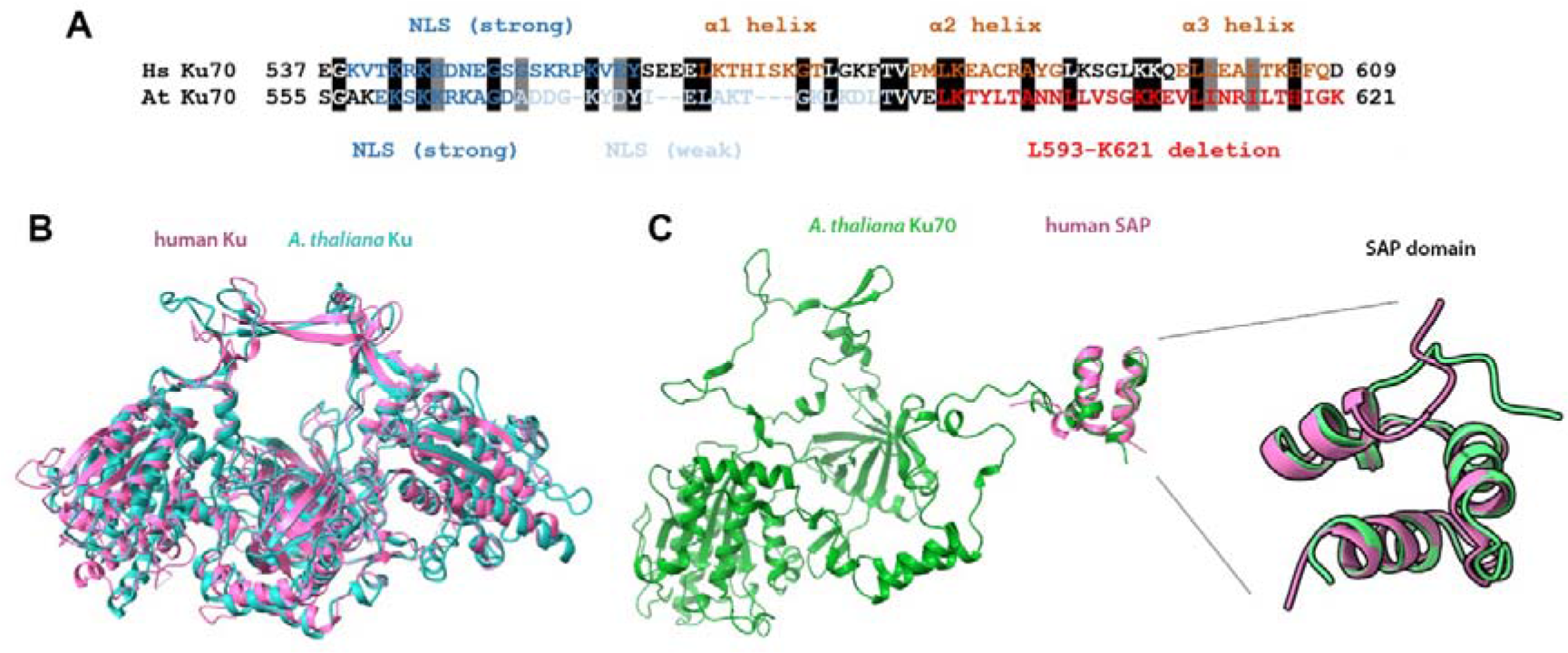
Structure of Arabidopsis Ku. (A) Sequence alignment of human (Hs) and Arabidopsis thaliana (At) Ku70 c-terminal region with predicted nuclear localization signals (NLS), alpha helices and depicted region which was deleted in ΔSAP variant. (B) Superimposed structures of human Ku obtained by X-ray crystalography (1JEQ; violet) and Arabidopsis Ku predicted by AlphaFold (cyan) to show the similarity of the core complexes (without Ku80 and Ku70 C-termini). (C) Structure of Arabidopsis Ku70 predicted by AlphaFold including the SAP domain and the flexible linker. The structure of human SAP domain determined by NMR (1JJR; violet) is superimposed over Arabidopsis SAP (green).

To analyze the impact of SAP domain on Ku-DNA binding, we generated Arabidopsis ΔSAP complex lacking the helix-extended loop-helix motif, but retaining the putative nuclear localization signal, by heterologous expression in insect cells (Figures 1A and 2A). Native polyacrylamide gel electrophoresis demonstrated that the ΔSAP complex forms stable heterodimers (Figure 2B). Effect of SAP on Ku-DNA interaction was assessed by electrophoretic mobility shift assay (EMSA) with DNA duplex probe that can accommodate up to four Ku complexes. Whereas wild type Ku readily formed complexes containing probe bound with four Ku molecules, higher order complexes form less efficiently with ΔSAP and products with four Ku were not detected in the tested concentration range (Figure 2C). This result collaborates data with human Ku and indicates the contribution of the SAP domain to DNA binding.

**Figure 2.**
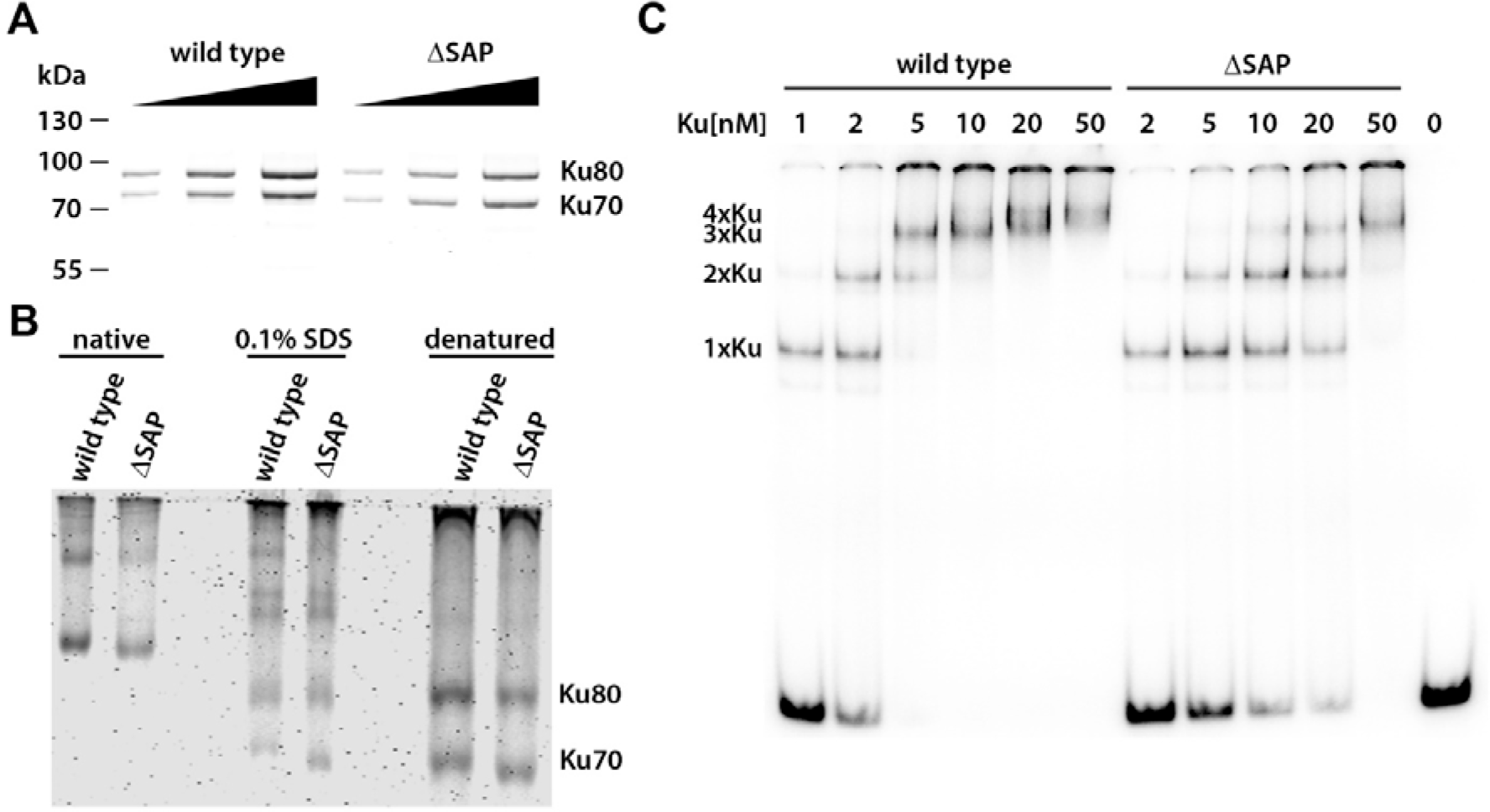
Biochemical characterization of the Ku ΔSAP complex (A) SDS-PAGE of purified recombinant Ku complexes stained by SimplyBlue SafeStain (B) Native-PAGE of recombinant Ku complexes demonstrates that deletion of Sap does not hinder dimerization. (C) EMSA of a radioactively labeled 98-bp DNA probe (0.3 nM) with increasing concentrations of recombinant Ku complexes. Positions of Ku-DNA complexes with different stoichiometries are indicated on the left side of the gel.

In our previous work, we have proposed that Ku-DNA association is a multistep process consisting of the initial interaction of DNA end with the Ku channel, Ku docking onto DNA, and sliding of Ku along DNA (Figure 3A) (12). To study the contribution of SAP to Ku-DNA binding more in detail, we used an assay based on protein induced fluorescence enhancement (mwPIFE) (33). In this assay, protein bound in immediate vicinity of a DNA-attached Cy3 fluorophore enhances emitted fluorescence, which can be used to measure protein-DNA interaction. We used DNA probes immobilized through one end to NeutrAvidin-coated microwell plates via biotin, while the opposite end was available for Ku binding. We have previously determined that the minimal binding site for Arabidopsis Ku is 13 bp (12). Therefore, we used 13 and 15 bp probes with Cy3 attached to DNA termini to assess the initial Ku-DNA interaction (Figure 3B). A 30 bp probe with Cy3 positioned 15 bp from DNA ends was used to monitor DNA docking, because in this setting PIFE should occur when Ku is fully loaded on DNA. And finally, Ku translocation along DNA was tested using a 55 bp probe with Cy3 placed 40 bp from free termini. All these probes showed decreased PIFE signal with ΔSAP compared to wild type Ku (Figure 3B). Nevertheless, the difference in the PIFE signal between wt and ΔSAP was much smaller with the 55 bp probe indicating that Ku sliding along DNA is less affected than the initial binding steps.

**Figure 3.**
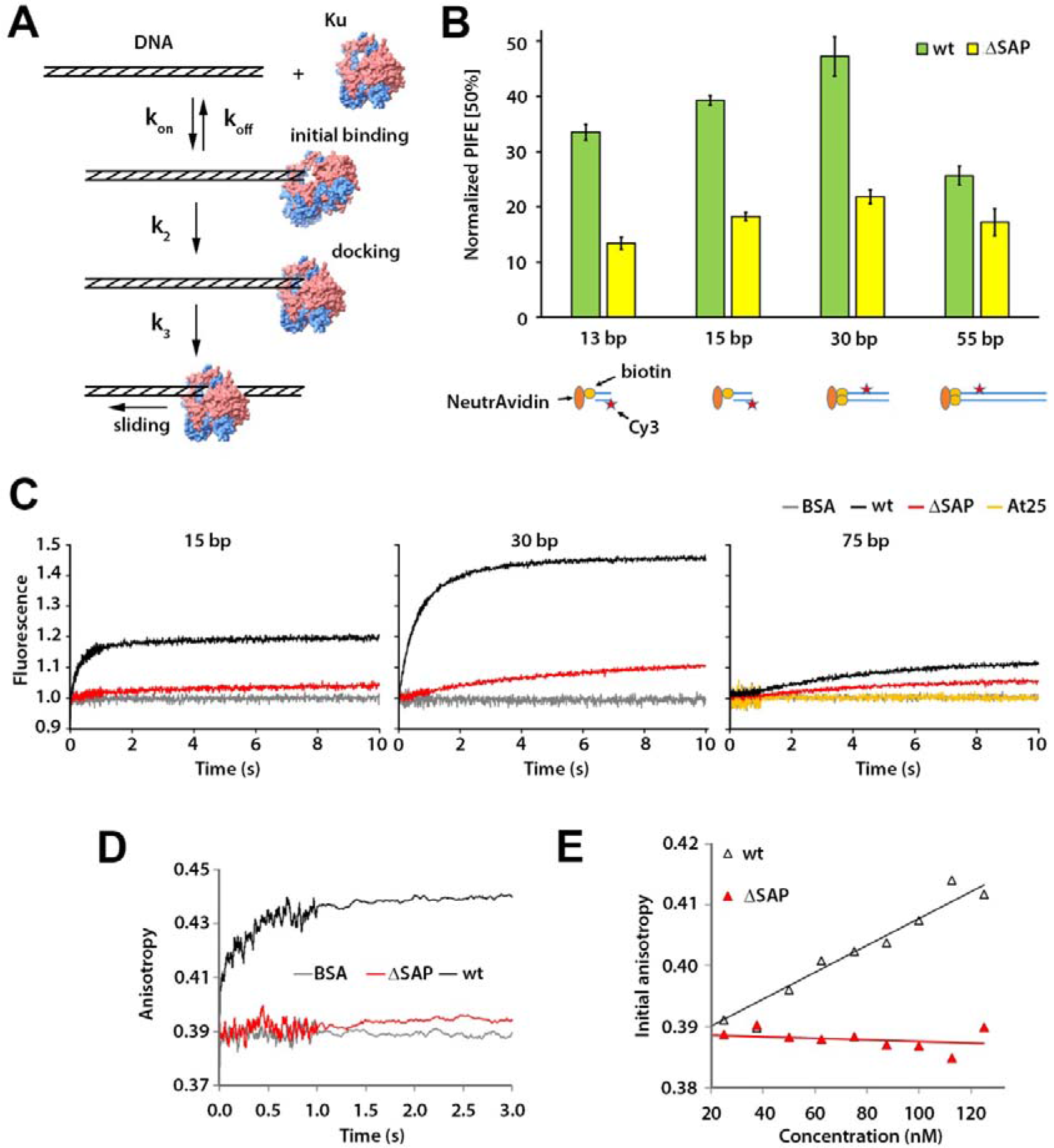
Mechanism of Ku DNA interaction. (A) Model of different phases of Ku DNA interaction. (B) Steady-state interaction of Ku at different position along the DNA probe measured by mwPIFE. PIFE was calculated as the relative difference in fluorescence upon protein binding. Error bars represent standard deviations from two replicas. (C) Kinetic analysis of Ku-DNA binding by stopped-flow PIFE using the 15-, 30-, and 75-bp DNA probes. Each trace represents the average of at least five individual measurements. The non-binding Ku variant (At25) was used as a control in the experiment with the 75 bp probe. (D) Anisotropy data from the stopped-flow measurements with the 30bp probe. (E) Chart showing the dependence of the initial anisotropy (time point zero) on the concentration of the Ku complex.

We next performed a kinetic analysis of Ku-DNA binding in stopped-flow system by scoring PIFE at millisecond intervals upon mixing Ku with DNA probes. We used a similar set of oligonucleotide probes as in the steady state measurements, with the exception that the 55 bp probe was extended to 75 bp, one end was blocked with biotin-neutravidin interaction, and the Cy3 fluorophore was placed 60 bp from the free end. Both wild type and ΔSAP produced PIFE signals that increased with time generating kinetic curves (Figures 3C and S2). The kinetic curves were measured for different protein/DNA molar ratios, and the concentration dependences of the observed rates were used to calculate the kinetic constants related to the individual steps of Ku-DNA interaction (Table 1, Figure S2).

**Table 1.** Kinetic parameters for DNA interaction with wild type and ΔSAP. Kinetics analysis was performed with three different DNA substrates to identify the effect of the SAP domain on individual kinetic phases of Ku-DNA interaction. The 15 bp and 30 bp probes show the fast initial binding followed by a docking phase, 15 bp was used to monitor the kinetics of the sliding phase. The estimates for the rate constants of Ku-DNA association (k_on_) and dissociation (k_off_), docking (k_2_) and sliding (k_3_) phases were obtained by analytical fitting from the concentration dependence of the observed rates (Figure S2). The reported equilibrium dissociation constant was calculated from the rate constants (K_D_ = k_off_/k_on_).

|  |  | *Initial binding* |  |  | *Docking* | *Sliding* |
| --- | --- | --- | --- | --- | --- | --- |
| | | $k_{on}$ | $k_{off}$ | $K_D$ | $k_2$ | $k_3$ |
| | | $\text{nM}^{-1} \cdot \text{s}^{-1}$ | $\text{s}^{-1}$ | $\text{nM}$ | $\text{s}^{-1}$ | $\text{s}^{-1}$ |
| wt | 15 bp | $0.021 \pm 0.005$ | $2.0 \pm 0.4$ | 95 | $0.6 \pm 0.1$ | - |
| ΔSAP | 15 bp | $0.002 \pm 0.001$ | $0.36 \pm 0.09$ | 180 | $0.2 \pm 0.1$ | - |
| wt | 30 bp | $0.023 \pm 0.005$ | $1.4 \pm 0.4$ | 61 | $0.78 \pm 0.07$ | - |
| ΔSAP | 30 bp | $0.002 \pm 0.001$ | $0.21 \pm 0.12$ | 105 | $0.09 \pm 0.02$ | - |
| wt | 75 bp | - | - | - | - | $0.19 \pm 0.05$ |
| ΔSAP | 75 bp | - | - | - | - | $0.14 \pm 0.03$ |

Ku interaction with 15 bp and 30 bp probes revealed two kinetic phases that reflect the fast initial interaction followed by slower conformational rearrangement, “docking” of the complex. The third single kinetic phase, corresponding to sliding, was monitored with the 75 bp probe. Deletion of the SAP domain substantially slowed the initial interaction and docking phases, but did not have a profound impact on sliding (Figure 3C, Table 1). Anisotropy measurements with the 30 bp probe confirmed the fast formation of the initial interaction complex in wild type, which was not observed with ΔSAP (Figure 3D,E). Together, these findings indicate that the SAP domain facilitates the initial Ku-DNA interaction, while imposing minimal impact on subsequent sliding phase.

Arabidopsis mutants lacking Ku are viable, but exhibit partial telomere deprotection and sensitivity to genotoxic agents (20,22). To assess the function of SAP in DNA repair and telomere protection, we generated transgenic Arabidopsis with disruption in the endogenous KU70 gene, carrying complementing KU70 constructs either with or without the SAP domain. Western blot analysis of several independent transgenic lines showed that the expression level of the Ku70 ΔSAP variant was substantially lower than the level of wild type Ku70 (Figure S4). The lower level of Ku70 ΔSAP does not appear to be caused by proteasome-mediated degradation and rather indicates a less efficient protein expression (Figure S4C).

Arabidopsis ku mutants have long telomeres due to unregulated extension by telomerase. Terminal restriction fragment analysis revealed that telomeres in *Δ*SAP plants were 3-6 kb, which is within the range of plants containing wild type Ku, whereas telomeres in Ku-deficient plants were over 10 kb (Figure 4A). Another characteristic of telomere deprotection in ku70 mutants is increased telomeric recombination, leading to the formation of extrachromosomal telomeric circular DNA molecules (t-circles) (25). The telomeric circle amplification assay showed that the level of t-circles in *Δ*SAP plants is comparable to wild type (Figure 4B). This data, together with telomere length analysis, indicates that despite its decreased DNA-end binding efficiency and lower expression in transgenic plants, Ku70 ΔSAP is fully proficient in telomere protection.

**Figure 4.**
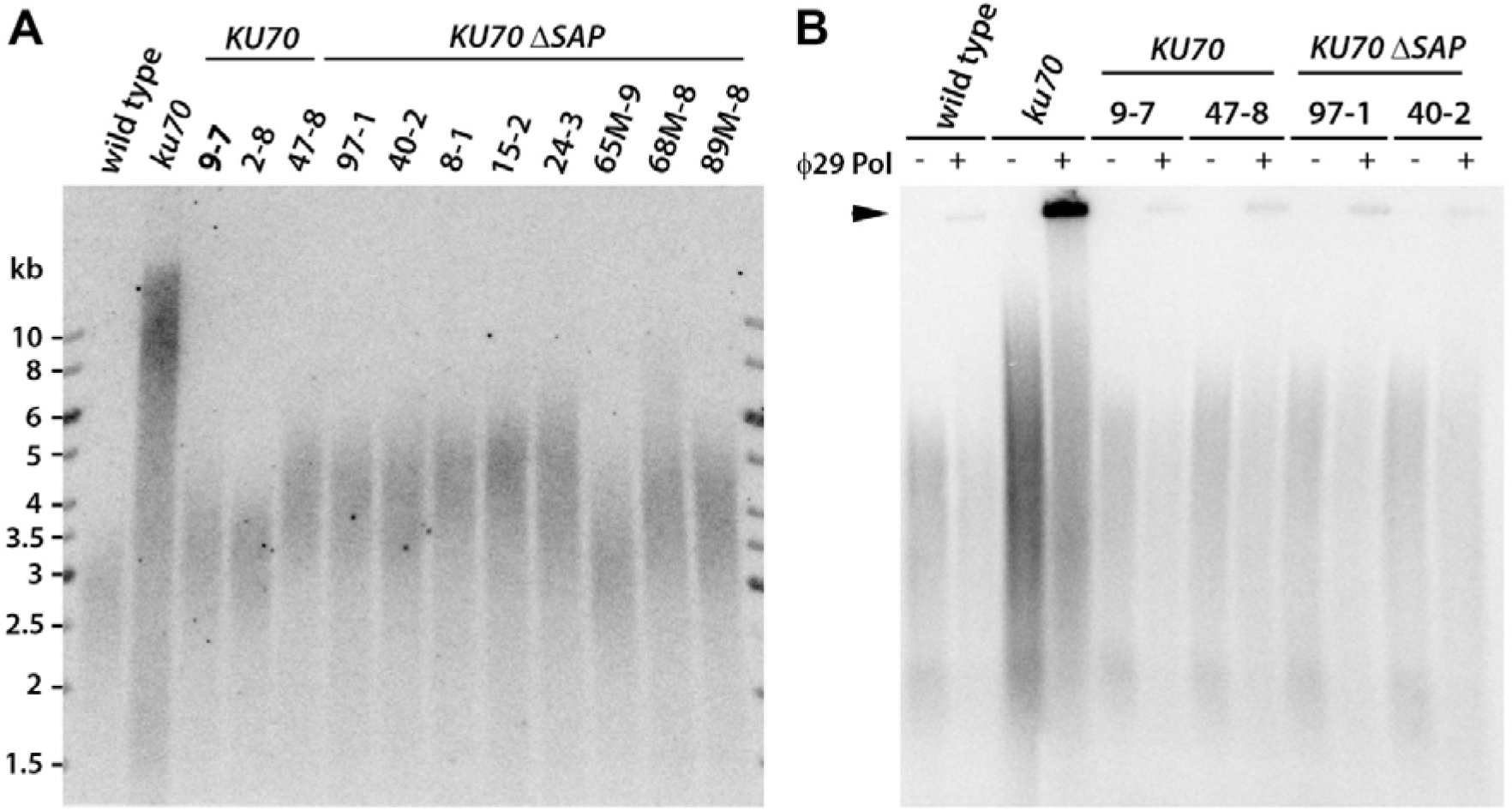
Effect of SAP on telomere maintenance. (A)Terminal restriction fragment length analysis with Tru1I-digested genomic DNA from wild type, ku70 mutants, and ku70 mutants complemented either with wild type KU70 or ΔSAP KU70 constructs. Several independent complemented lines in T2 generation were analyzed. (B) The presence of t-circles detected by t-circle amplification assay. Reactions without phi29 polymerase were run in parallel as a control. The signal from t-circles is indicated by the arrowhead. Two independent lines were analyzed for each complementation construct.

We next tested the proficiency of *Δ*SAP plants in DNA repair. Arabidopsis ku70 and ku80 mutants are highly sensitive to the radiomimetic drug bleomycin (12,34). The majority of seedlings of ku70 mutants germinated on agar plates supplemented with a low concentration of bleomycin (50 ng/mL) either do not develop or have malformed true leaves, whereas wild type plants exhibit similar aberrations at three times higher concentration (Figure 5A). This sensitivity is also apparent in the root growth assay, whereby root development in ku70 mutants is completely arrested at agar plats supplemented with 25 ng/mL of bleomycin, whereas roots in wild type develop normally (Figure 5B). Intriguingly, *Δ*SAP plants behave as wild type in both seedling germination and root growth assays and do not exhibit increased sensitivity to bleomycin (Figure 5). This data indicates that the Ku complex lacking the SAP domain is not substantially hindered in DNA repair.

**Figure 5.**
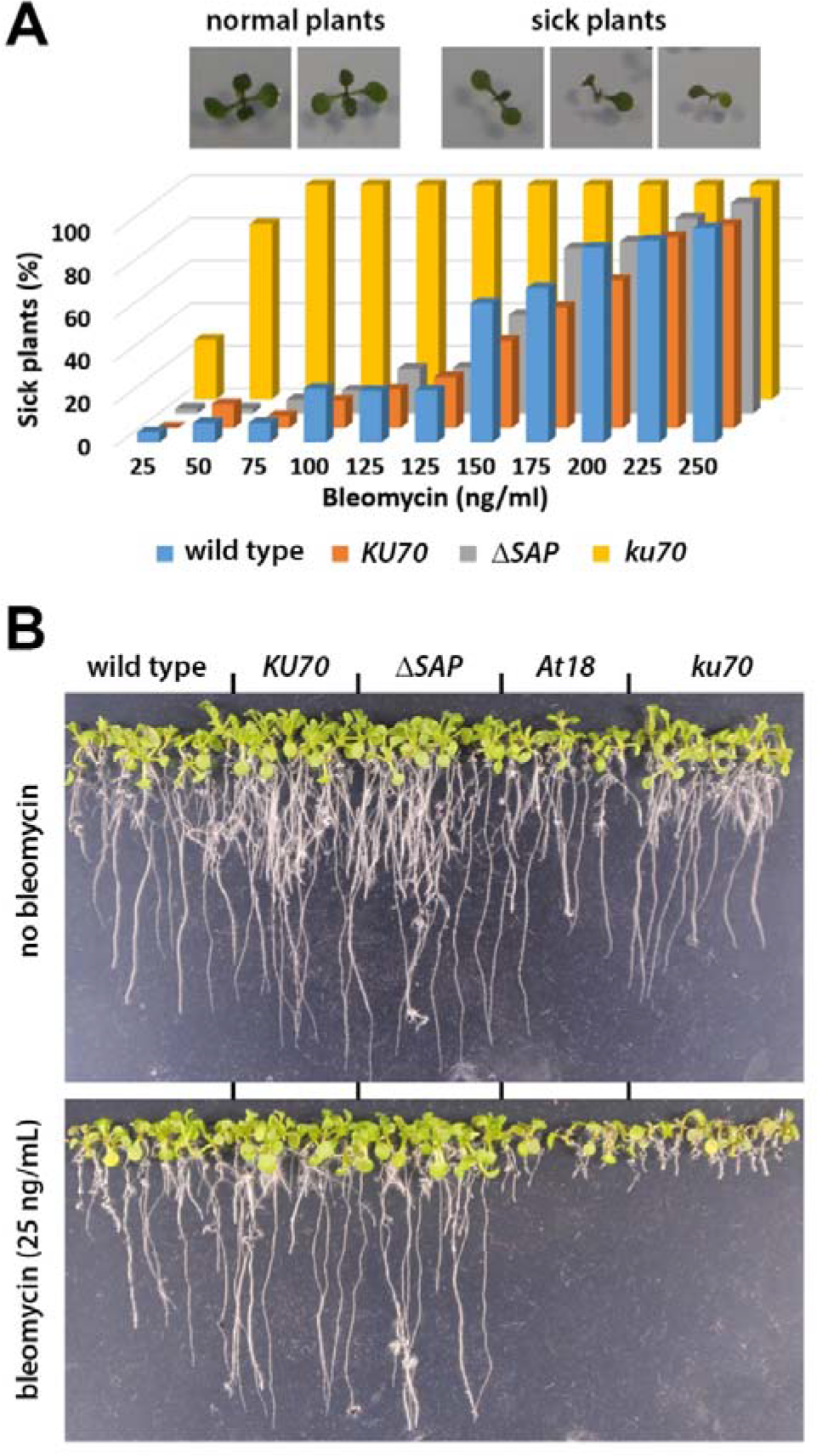
Sensitivity to bleomycin. (A) Seedling growth assay. Pictures show normally developed and sick seedlings with malformed true leaves 11 days after germination on media supplemented with bleomycin. 3D column chart indicates frequency of normal and sick seedlings germinated on media with different concentrations of bleomycin. At least 50 seedlings were count for each category. (B) Root growth on media with or without bleomycin 11 days after germination.

c-NHEJ represents a primary route of DSB repair in G0/G1 and is determined by the binding of Ku to broken DNA ends, which limits excessive end resection. In the absence of Ku, DSBs can be repaired by alt-EJ, which can be distinguished by producing larger deletions, typically more than 10 bp, at the site of repair (35,36). To evaluate whether the less efficient binding of Ku70 ΔSAP to DNA ends skews DSB repair towards alt-EJ, we designed an end joining assay based on ligation of a linear DNA fragment transfected into plant protoplasts (Figure S4). End joining products were PCR amplified 24 hrs after transfection using primers spanning 225 bp of the ligation site, subjected to next generation sequencing, and the joined ends were analyzed to determine the extend for DNA resection prior to ligation. Surprisingly, a large portion of joined ends exhibited deletions larger than 10 bp, even in Ku proficient protoplasts (Figure 6). We suspect that this reflects rapid nucleolytic resection of naked chromatin-free DNA prior to its association with Ku, rather than predominance alt-EJ in Arabidopsis protoplasts. Nevertheless, the fraction of NHEJ events containing intact ends or deletions smaller than 10 bp was substantially larger in Ku-proficient cells compared to protoplasts from ku70 mutants. We also included in the analysis protoplasts from At18 plants that contain a Ku complex that is hindered in sliding along DNA (12). We anticipated that the entrapment of Ku at DNA ends due to inefficient sliding would provide better end-protection and less resection. Indeed, the fraction of end-joining products with intact or less resected ends was greater in At18 than in plants with wild type Ku. *Δ*SAP protoplasts exhibited a similar end-resection pattern as Ku-proficient plants, suggesting that deletion of the SAP domain does not substantially compromise c-NHEJ.

**Figure 6.**
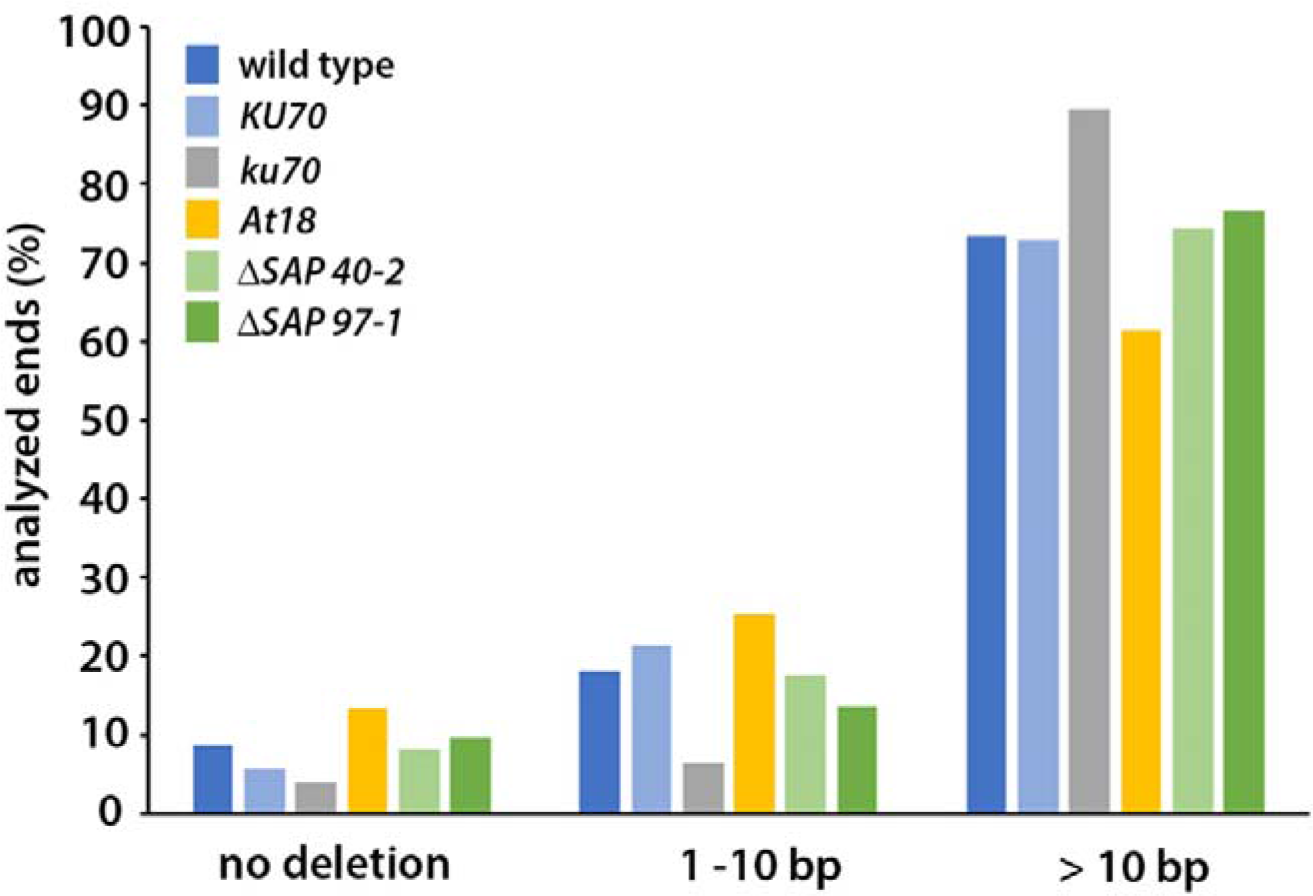
Sequence analysis of fusion junctions in the end-joining assay. Frequency of DNA ends with different degree of resection scored in the end joining assay. The extent of the resection was calculated from the fusion point. Each ligated product provided information on two DNA ends. More than 120,000 amplicons were scored in each category.

## Discussion

Ku plays a crucial role as an early responder to DNA damage by rapidly binding to DNA double-strand breaks (DSBs) and other free ends of DNA duplexes. The atomic structure of Ku provided crucial insight into the mechanism behind its sequence non-specific affinity for DNA ends (9). The Ku complex adopts a ring-like structure with a cradle at its base that allows DNA to thread through a preformed channel, thereby enabling Ku to load exclusively through DNA ends. The inner surface of the channel is composed of positively charged amino acid residues, which can potentially attract the negatively charged DNA backbone. Notably, the channel aperture appears smaller than the diameter of the duplex DNA, and its inner lining resembles a helical thread, suggesting that Ku may bind to DNA akin to a bolt on a nut.

Such a mode of interaction predicts that loading requires precise positioning of DNA end at the entry to the Ku channel and may represent the rate-limiting step. Our stopped-flow analysis indicates that Ku-DNA binding occurs in two kinetic phases, which we interpret as rapid initial contacts between Ku and DNA, followed by a slower step that may represent docking Ku onto DNA analogous to screwing a nut onto a bolt (Figure 3A) (12). Once Ku is loaded, it can freely slide along DNA. The positioning of the DNA end is likely driven by electrostatic interactions with positively charged amino acids in the entry of the channel and the cradle facing the DNA. This is supported by the observations that mutations reverting the charge at these positions abolish Ku-DNA binding (11,12).

Studies on human Ku (15,37) together with our data in Arabidopsis indicate an evolutionarily conserved role of the Ku70 SAP domain in DNA binding. Our work implies that the SAP domain is involved in the initial steps of Ku-DNA interaction. Kinetic analysis showed that the ΔSAP Ku-complex is impaired already in the first binding phase, suggesting its function in mediating first contacts with DNA or in stabilizing these initial interactions. Structure-based modeling predicted that positively charged patches on the surface of the SAP domain and the flexible linker may interact with DNA (15,16). Thus, the Ku70 C-terminus may act as an extended flexible holder that can grab free DNA end and drag it to the channel, or stabilize the initial interactions during loading of Ku onto DNA. This is further supported by the observation that ΔSAP variant of human Ku very rapidly dissociates from a DNA substrate in a FRET-based assay (37). This model is also consistent with a recent structural analysis of human Ku showing that in the DNA-free conformation, SAP is located at the leading face of the complex in the vicinity of the channel (17). This may also contribute to the directionality of Ku-DNA binding.

SAP could also potentially play a role in later steps of Ku-DNA interaction. For example, it could assist in Ku translocation along DNA by diffusion-ratchet mechanisms (38), or stabilize established interaction by preventing Ku from sliding off the DNA. However, we did not observe an impact of SAP on the kinetics of Ku translocation along DNA, and SAP appears to be positioned away from DNA once Ku is stably bound (17). Additionally, SAP does not affect Ku retention on DNA when it is in complex with DNA-PKcs, which displaces Ku from the DNA end to more internal positions (37,39). This supports the notion that SAP is required during the initial binding, but is dispensable for the stable Ku-DNA interaction of the fully loaded complex.

Curiously, despite reduced efficiency of DNA binding and lower expression, ΔSAP Ku appears to be proficient in telomere maintenance and c-NHEJ in Arabidopsis. In c-NHEJ, Ku binding to a DSB prevents excessive nucleolytic resection and coordinates processing activities to form ligatable ends. In the absence of Ku, DNA ends are exposed to DNA resection, which can reveal microhomologies between complementary strands on the opposite sites of a DSB. These sequence microhomologies are utilized by alt-EJ pathways to facilitate synapsis and ligation of broken DNA ends (3). Ku-deficient Arabidopsis exhibits sensitivity to bleomycin (12,34), and sequence analysis of EJ- junctions, generated either by re-ligation of CRISPR/Cas9 induced breaks or chromosome end-to-end fusions, revealed larger deletions and more frequent microhomologies typical for alt-EJ (36,40).

We found that ΔSAP Ku fully rescues bleomycin sensitivity of ku70 mutants. Furthermore, end-joining assay performed by transfecting naked linear DNA into mesophyll protoplast combined with high-throughput sequencing of fusion junctions showed a re-ligation pattern similar to wild type rather than to ku70 mutants in terms of frequency larger (>10 bp) deletions. This indicates that even the less efficient loading of ΔSAP Ku onto DSBs is still sufficient to support c-NHEJ and prevent access of competing nucleases. This also appears to be the case at telomeres. In Arabidopsis, Ku acts in protecting telomeres through their physical sequestration within the binding channel (12,23). This prevents the resection of chromosome ends replicated by the leading strand mechanism, resulting in blunt-ended telomeres. In the absence of Ku, telomeres are resected by EXO1 and other nucleases, exposing long 3’overhangs that serve as substrates for homologous recombination or extension by telomerase (23–25). Efficient protection of telomeres from excessive elongation by telomerase and homologous recombination in ΔSAP plants argues that, like in the context DSBs, the speed of Ku loading onto telomeres is not the decisive factor for outcompeting these other end processing activities.

## Acknowledgements

The project was supported by the Czech Science Foundation (19-21961S) to J.F. Scientific data presented in this paper were obtained with the support of the following Core Facilities (CF) of CEITEC Masaryk University: Plant Science CF, CF Bioinformatics, Genomics CF (supported by MEYS CR infrastructure project LM2023067) and Biomolecular Interactions and Crystallography CF of CIISB (supported by MEYS CR infrastructure project LM2023042 and European Regional Development Fund Project “UP CIISB” No. CZ.02.1.01/0.0/0.0/18_046/0015974). Z.P. was supported by the European Union’s Horizon 2020 research and Innovation program TEAMING 857560, the Czech Ministry of Education, Youth and Sports TEAMING CZ.02.1.01/0.0/0.0/17_043/0009632), ESFRI RECETOX RI (LM2023069) and EXCELES Onco LX22NPO5102.

## Supplementary information

**Figure S1.**
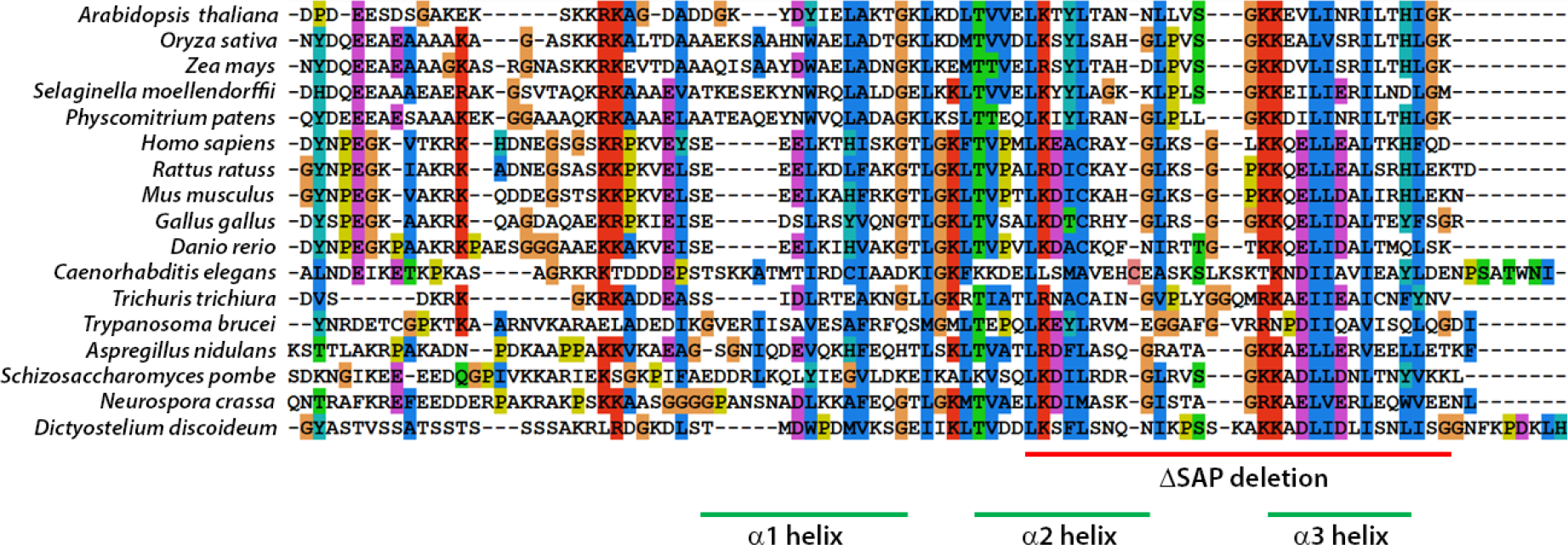
Multiple amino acid sequence alignment of Ku70 SAP domains from species representing different phylogenetic classes of eukaryotes. Regions spanning the predicted α-helixes (green lines) and the ΔSAP deletion analyzed in this study (red line) are indicated.

**Figure S2.**
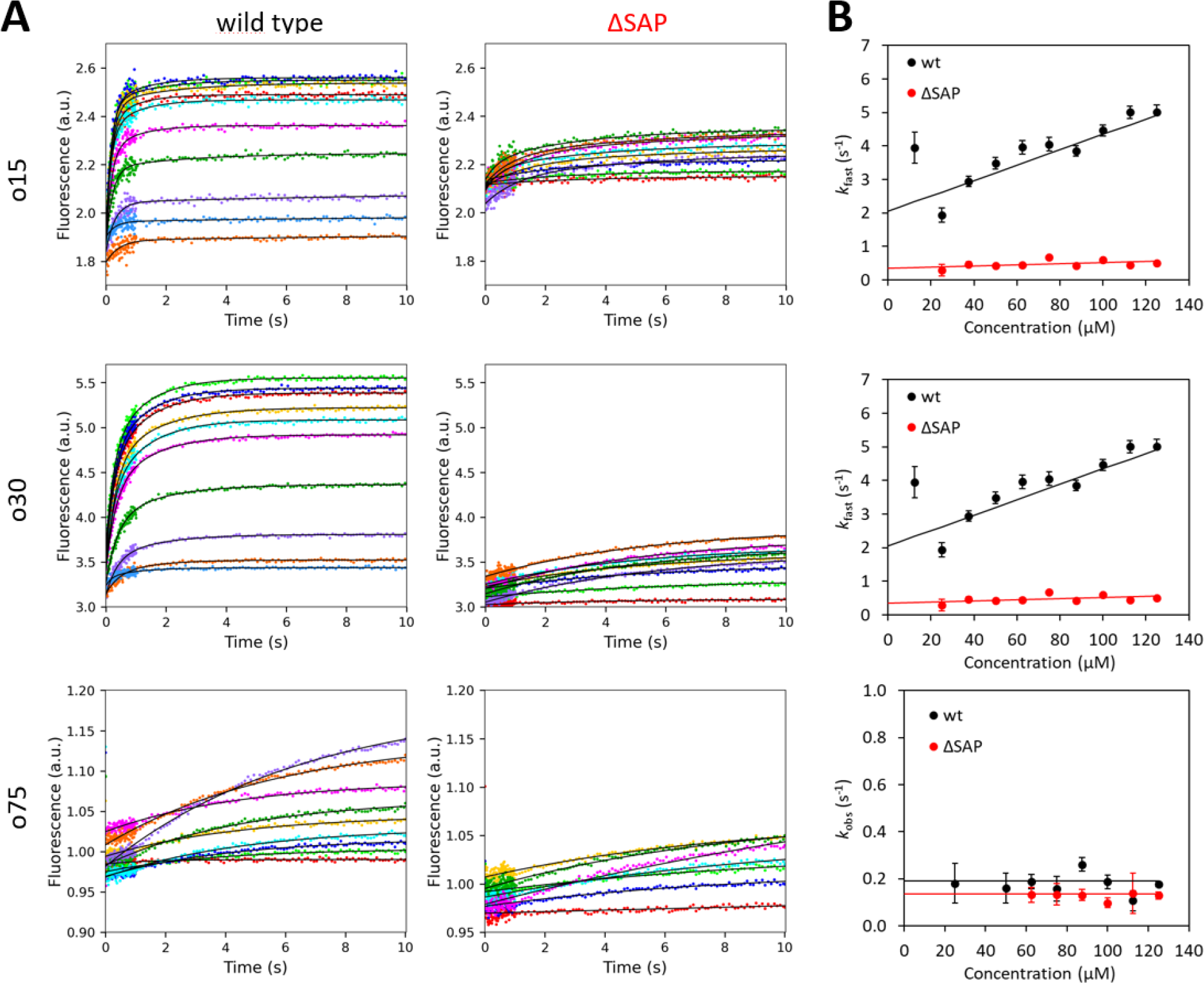
Analytical fitting of kinetic data. (**A**) The stopped-flow fluorescence traces (PIFE signal) were recorded upon mixing 12.5 nM ssDNA with 0 – 125 nM Ku. Each stopped-flow trace represents the average of at least five replicates. The solid lines represent fit to a single exponential function *f* = *F*_0_ + *A*_1_. (1 − *e*^−*k*^_*obs*_.^*t*^) for reaction with o75 or a double exponential function *f* = *F*_0_ + *A*_1_. (1 −*e*^−*k*^_*fast*_.^*t*^) + *A*_2_. (1 − *e*^−*k*^_*slow*_.^*t*^) for o15 and o30. It is important to note that the range of the y-axis is consistently fixed for graphs depicting both wild-type and ΔSAP data to ensure a fair comparison. (**B**) The concentration dependence of the observed rates of the fast initial phase derived from fitting o15 (upper graph) and o30 (middle graph) data in (A) provides the rate constants for initial association (*k*_on_) and dissociation steps (*k*_off_). The concentration dependence of the observed rates of the slow phase was used to get estimates for the rate constant of the docking phase (*k*_2_). The observed rates derived from single exponential fitting o75 data (bottom graph) provided an estimate for the first-order rate constant of the sliding phase (*k*_3_). The solid lines in the figure depict the linear model fitted to the data. The error bars in (B) represent the standard errors of the rate parameter estimates obtained through the exponential fitting.

**Figure S3.**
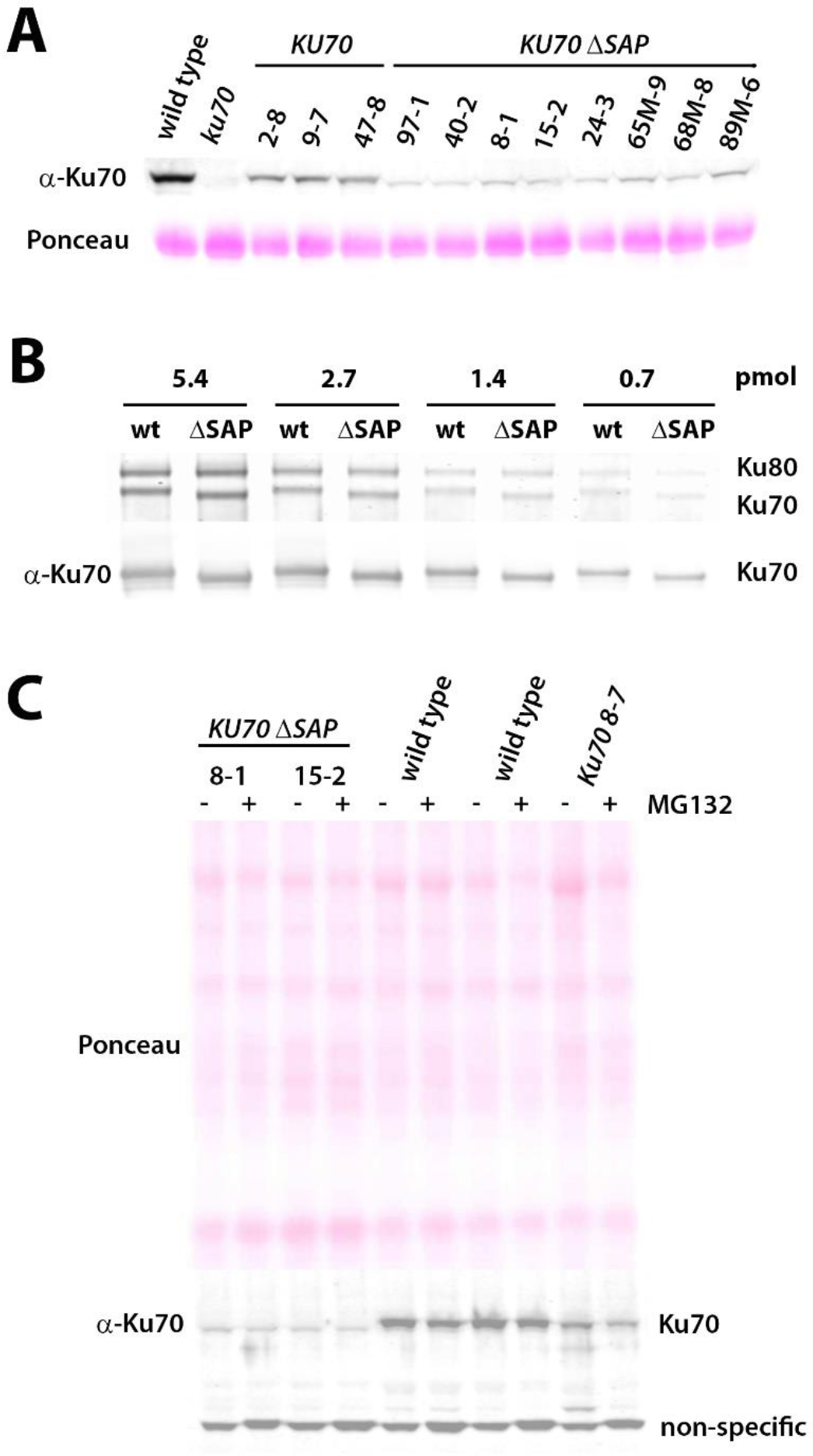
Ku70 expression in Arabidopsis complemented lines. (**A**) Protein levels of Ku70 in individual complemented lines anlyzed by SDS-PAGE and western blot immunodetection using antibodies raised against Arabidopsis Ku70. Ponceau S stained RuBisCo at the same membrane shows protein loadings. (**B**) Assay to determine whether the wild type and ΔSAP Ku70 proteins are equally recognized by the custom-made Ku70 antibody. Concentration range of purified recombinant Ku complexes was separated by SDS-PAGE and detected by SimplyBlue SafeStain (top panel). This was followed by western blot analysis using custom made Ku70 antibody. No difference in the signal intensity was detected. (**C**) Proteasome inhibition assay to determine whether the lower expression of ΔSAP Ku is due to degradation by a proteasome. Seedling were treated with the proteasome inhibitor MG132, extracted protein was separated on SDS-PAGE, and the Ku70 protein levels were analyzed by western blot immunodetection. Upper panel shows Ponceau S stained membrane with the total protein, and the lower panel indicates Ku70 signal detected by immunostaining with Ku70 antibody. No difference is apparent between MG132 treated and non-treated samples suggesting that lower level of ΔSAP Ku70 is caused by its proteasome degradation.

**Figure S4.**
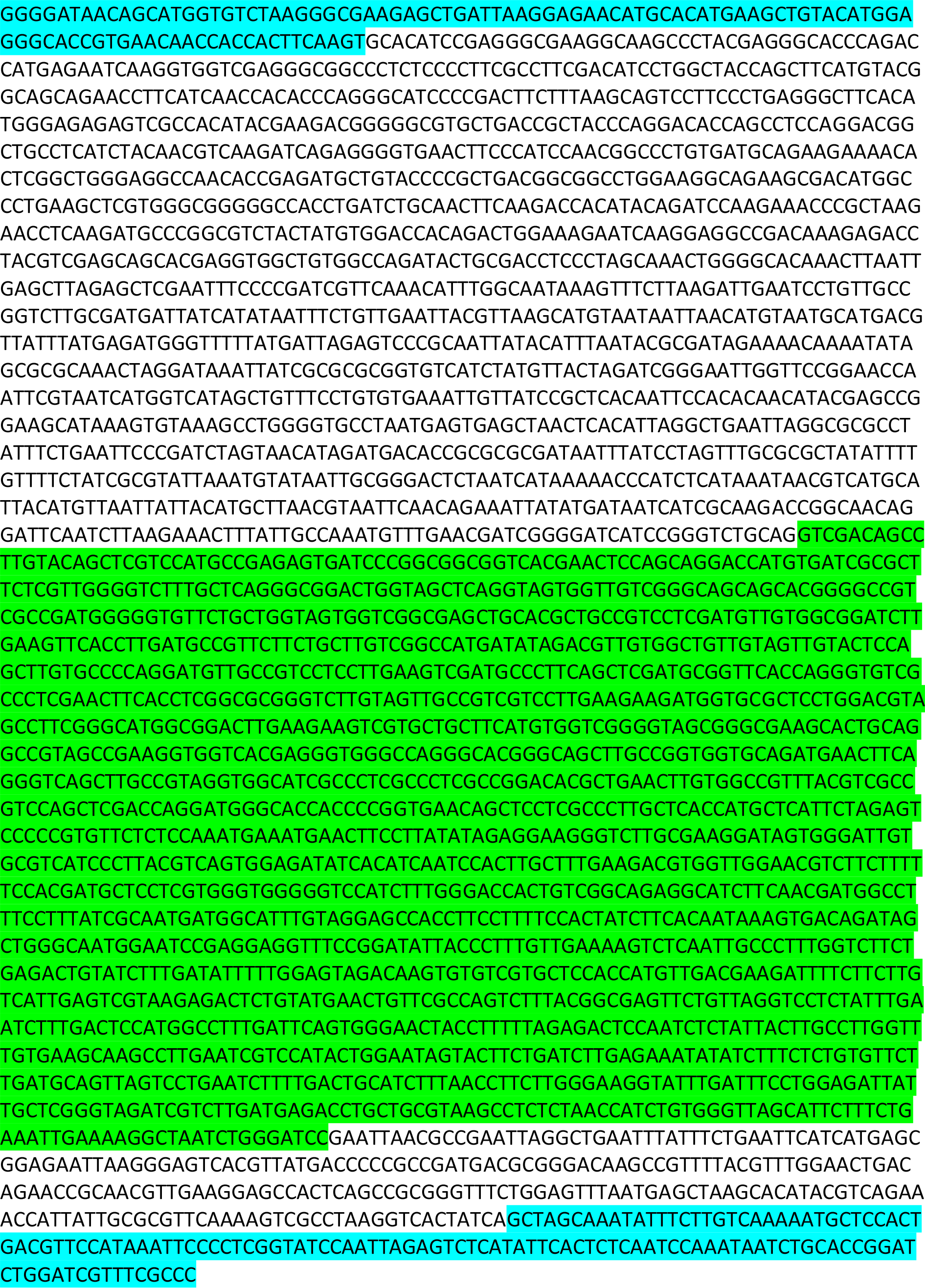
Sequence of the linear DNA fragment used in the end joining assay. The sequence spanning the fusion junction after fragment ligation in protoplasts that was PCR analyzed by Illumina sequencing is highlighted in blue. Region coding for 35S:GFP reporter is indicated in green.

